# ThermoFusion: A Multimodal Deep Learning Framework for Generalizable Prediction of Enzyme Thermostability

**DOI:** 10.64898/2026.07.04.736494

**Authors:** Yao Wei, Ivano Eberini, Fabian Meyer

## Abstract

Protein thermostability is a critical property for both industrial and biomedical enzyme applications, yet experimental evaluation of mutation-induced stability changes remains laborious and costly. Here, we present ThermoFusion, a hybrid deep learning framework that integrates 3D protein structure embeddings from ThermoMPNN with sequence-based embeddings from the pretrained protein language model ESM2 to predict the effects of single-point mutations on protein stability (ΔΔ*G*). ThermoFusion exhibits robust generalization, maintaining high predictive accuracy across out of distribution sequences with low identity to the training set – a scenario where many other machine learning models, including ThermoMPNN and state-of-the-art tools, perform poorly due to reliance on memorization. Benchmarking on a curated enzyme dataset comprising of 105 enzymes and 3144 mutations shows that ThermoFusion reliably identifies stabilizing mutations while accurately predicting stability for enzymes beyond its training set. These results establish ThermoFusion as a powerful tool for rational enzyme design beyond its training set.

## Introduction

Proteins are the essential macromolecules of life that carry out nearly all biological processes within cells. Composed of 20 canonical amino acids, the amino acid sequence encodes the three-dimensional (3D) structure of a protein, which in turn governs its stability, catalytic activity, and interaction networks. Understanding the relationship between sequence, structure, and stability is therefore fundamental for molecular biology, biotechnology, and medicine (2, 24).

Protein stability refers to a protein’s ability to maintain its native folded conformation under specific physiological conditions (e.g., temperature, pH). Thermodynamically, stability is governed by the balance of non-covalent interactions – including hydrogen bonds, ionic interactions, and hydrophobic forces (16). It can be quantified using parameters such as folding free energy (Δ*G*) (5), heat capacity change (Δ*Cp*) (32), or melting temperature (Tm) (37). In the context of protein engineering, the change in folding free energy upon mutation (ΔΔ*G*) is particularly informative, as it directly quantifies the effect of an amino acid substitution on protein stability. By comparing the folding free energies of wild-type and mutant proteins, ΔΔ*G* provides a practical metric to guide the design of stabilizing or destabilizing mutations (20).

A major goal of protein engineering is to design proteins with increased stability and enhanced functionality for scientific, industrial, and biomedical applications (21, 25, 33). Approaches such as directed evolution, pioneered by Frances H. Arnold and colleagues (3), emulate natural evolutionary processes through iterative rounds of mutation and selection to obtain improved variants. While effective, experimental strategies require the construction and screening of large mutant libraries, making them time-consuming and costly. Nevertheless, mutagenesis is widely used in protein engineering because it can improve enzyme functionality and provide interpretable insights into how sequence changes affect protein structure, stability, and function (26). While accurately predicting the impact of mutations on protein stability, could substantially reduce experimental burden and guide rational protein design.

Experimental methods such as differential scanning calorimetry (DSC), circular dichroism spectroscopy, and fluorescence spectroscopy provide detailed thermodynamic measurements for individual protein sequences, but their low throughput and substantial resource demands limit their ability to probe protein stability across large regions of sequence space (4). Physicochemical-based computational approaches, such as molecular dynamics simulations (12), MM/GBSA calculations (6) and free energy perturbation (35), offer detailed mechanistic insights but remain computationally expensive for large-scale mutation screening. Empirical energy-based tools, such as Rosetta (17) and FoldX (9), improve efficiency but are limited by their parameterization and generalization capacity. Recent advances in machine learning (ML) have improved protein engineering by enabling data-driven prediction of sequence - structure - function relationship (13). Classical ML algorithms – including random forests, support vector machines, and gradient boosting methods – perform effectively on small datasets with handcrafted features. More recently, deep learning architectures, such as convolutional neural networks, and Transformer-based protein language models, have demonstrated superior performance by automatically learning rich sequence and structural representations (15, 42). In particular, large-scale pretrained models, such as ESM2, capture evolutionary and biophysical information directly from sequence data, significantly improving downstream stability prediction tasks (43).

In this work, we propose ThermoFusion, a deep learning framework for predicting single-mutation-induced protein stability changes. ThermoFusion integrates complementary information from both protein structure and sequence by combining structural embeddings derived from ThermoMPNN with sequence embeddings extracted from the pretrained protein language model ESM2. By fusing structural and sequence-level representations, ThermoFusion leverages the strengths of both modalities to achieve improved accuracy and robustness in protein thermostability prediction. Importantly, ThermoFusion exhibits strong generalization on enzymes that differ from its training set, outperforming other machine learning models that are often unable to accurately predict the unseen proteins due to reliance on memorization rather than true understanding of underlying structural and sequence relationships. This highlights ThermoFusion’s ability to capture fundamental biophysical principles, making it broadly applicable to diverse protein families and mutation contexts.

## Results

### A. Model Architecture Overview

To predict the protein single-mutation-induced thermostability changes (ΔΔ*G*), we developed ThermoFusion (**Figure** 1), a hybrid sequence-structure deep learning framework that integrates geometry-aware structural representations with large-scale pretrained sequence embeddings. ThermoFusion builds on ThermoMPNN (10) and adopts its transfer-learning strategy to obtain structure-conditioned residue embeddings derived from a ProteinMPNN (7) backbone, thereby encoding three-dimensional (3D) geometric context from protein structures. In parallel, contextual residue-level sequence embeddings are extracted from the pretrained ESM2 model (esm2_t33_650M_UR500) (19), enabling the model to capture evolutionary signals and sequence-derived biochemical constraints.

**Fig. 1.**
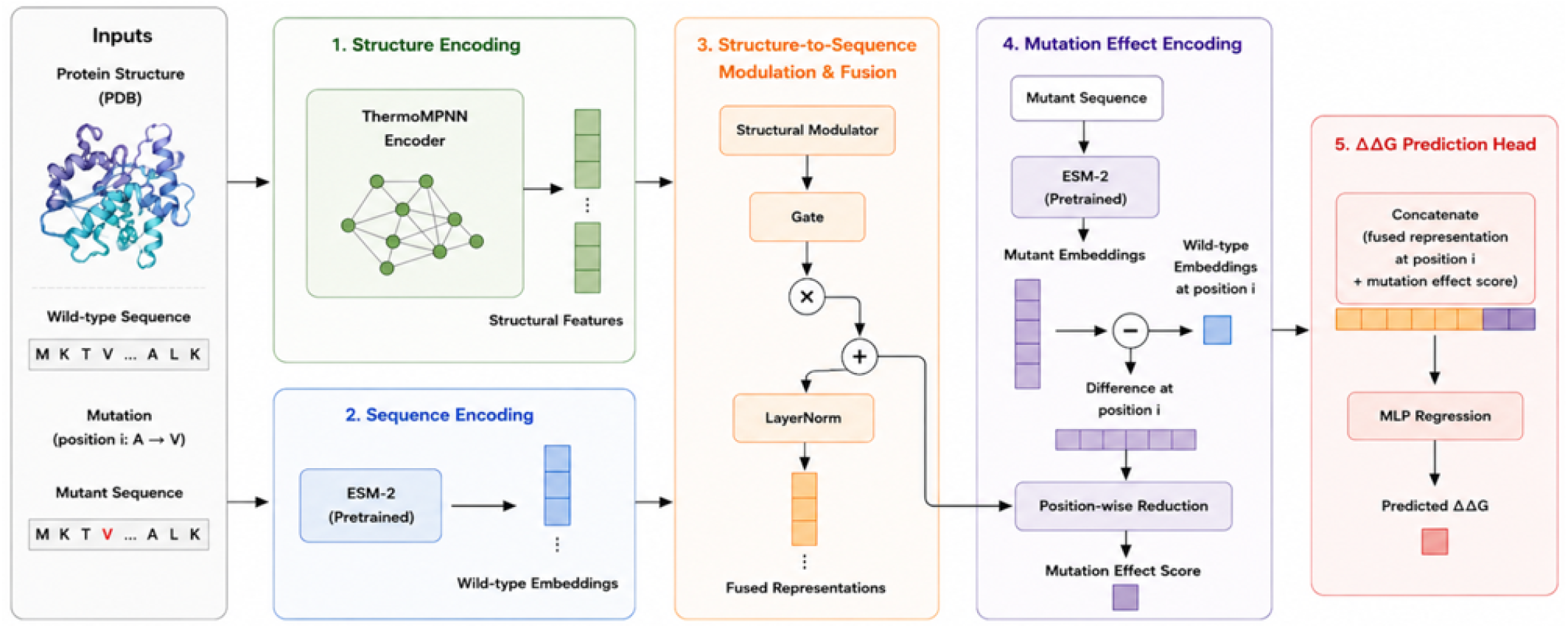
Overview of ThermoFusion Architecture. Sequence embeddings are extracted using the pretrained ESM2 model, while structure-aware residue graph representations are generated by ThermoMPNN. The resulting representations are aggregated and passed to a MLP to predict folding free energy change (ΔΔ*G*) between wild-type and mutant proteins

The ThermoMPNN-derived structural features are projected into the ESM2 embedding space through a multilayer perceptron (MLP) and then passed through a learnable gating module that regulates the contribution of structural information at each residue position. The gated structural representation is fused with the corresponding wild-type ESM2 residue embedding using residual addition followed by layer normalization, producing unified structure-aware sequence representations.

Given a protein structure and a mutation *m* = (*i, a*_*wt*_ →*a*_*mut*_), where *i* denotes the mutated residue position, the model predicts the thermodynamic stability change:

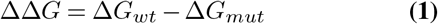

where negative values indicate destabilizing mutations and positive values indicate stabilizing mutations.

To capture mutation-induced perturbations, mutant sequence embeddings are recomputed using ESM2, and the difference between mutant and wild-type embeddings at the mutated residue is used as a mutation effect representation. This difference vector is compressed through a lightweight position-wise MLP to generate a scalar mutation effect score. Finally, the fused structure-aware residue embedding at the mutated position is concatenated with this mutation effect score and passed through a MLP to produce the final numerical (ΔΔ*G*) prediction.

### B. Comparison of ThermoFusion and Baseline ThermoMPNN Across Diverse Datasets

To evaluate the effectiveness of ThermoFusion, we compared its prediction performance against the baseline models, ThermoMPNN, across five published datasets: FireProt (HF) (10), S669 (27), S2648 (8), Q3421 (29), and ThermoMutDB (41). Only single-point mutations with experimentally measured ΔΔ*G* values were included in the evaluation. Dataset statistics, including each dataset’s sequence identity relative to the Megascale and FireProt training sets, are summarized in **Table** 1.

**Table 1.** Datasets Information for Model Comparison Study.

| Dataset | N (PDBs) | N (Mutations) | Sequence identity |  |  |
| --- | --- | --- | --- | --- | --- |
|  |  |  | ≤30% | 30–60% | >60% |
| FireProt (HF) | 80 | 2471 | – | 47.5% | 52.5% |
| S669 | 88 | 669 | – | 99.6% | 0.4% |
| S2648 | 351 | 2468 | 6.0% | 47.0% | 47.0% |
| Q3421 | 382 | 3421 | 0.3% | 53.1% | 46.6% |
| ThermoMutDB | 268 | 3389 | 0.4% | 85.8% | 13.8% |

The comparative study between ThermoFusion and ThermoMPNN is summarized in **Table** 2. Overall, ThermoFusion consistently outperformed ThermoMPNN across all benchmark datasets, exhibiting substantial reductions in root mean squared error (RMSE) together with notable improvements in Spearman’s correlation coefficient (SCC) and Pearson’s correlation coefficient (PCC). The largest performance gains were observed on FireProt (HF), where ThermoFusion improved SCC from 0.638 to 0.836 and PCC from 0.633 to 0.840 while reducing RMSE from 1.527 to 1.034. Similar improvements were also observed for the protein with sequence identities below 40% relative to the Megascale and FireProt training sets, indicating that ThermoFusion generalizes more effectively to proteins with limited sequence similarity to the training data.

**Table 2.** Performance comparison of ThermoFusion and ThermoMPNN across benchmark datasets.

| Dataset | Method | RMSE | SCC | PCC |
| --- | --- | --- | --- | --- |
| FireProt (HF) | ThermoMPNN | 1.527 | 0.638 | 0.633 |
|  | **ThermoFusion** | **1.034** | **0.836** | **0.840** |
| FireProt (HF) ≤40% | ThermoMPNN | 1.698 | 0.622 | 0.631 |
|  | **ThermoFusion** | **1.361** | **0.699** | **0.746** |
| S669 | ThermoMPNN | 1.530 | 0.476 | 0.419 |
|  | **ThermoFusion** | **1.470** | **0.492** | **0.491** |
| S2648 | ThermoMPNN | 1.175 | 0.639 | 0.625 |
|  | **ThermoFusion** | **0.956** | **0.767** | **0.776** |
| Q3421 | ThermoMPNN | 1.698 | 0.598 | 0.544 |
|  | **ThermoFusion** | **1.518** | **0.709** | **0.647** |
| ThermoMutDB | ThermoMPNN | 2.010 | 0.662 | 0.628 |
|  | **ThermoFusion** | **1.698** | **0.739** | **0.716** |

ThermoFusion also demonstrated consistent improvements on larger and more diverse datasets, including S2648, Q3421, and ThermoMutDB, suggesting enhanced robustness across diverse protein families. Although the improvements on S669 were comparatively smaller, ThermoFusion still achieved better overall correlation metrics than ThermoMPNN. These improvements highlight the importance of integrating both modalities for robust generalization.

### C. Benchmarking Different Prediction Models on the Curated Enzyme Dataset

Enzymes are invaluable biocatalysts with essential roles in numerous industrial applications, including biofuel production, food processing, detergent formulation, and pharmaceutical manufacturing. Thermostability is an essential attribute in enzyme engineering, as it has a direct impact on both enzyme performance and longevity under operational conditions. To evaluate ThermoFusion’s performance on enzymes, we curated a dataset of single-point enzyme mutations from the five datasets used in the above baseline model comparison study. After processing, the curated dataset comprises 105 enzymes spanning five major enzyme classes (Supplementary Table S1), with a total of 3,144 mutation sites. We subsequently benchmarked ThermoFusion against 12 widely used protein mutation thermostability prediction tools, including 6 machine learning-based and 6 non-machine learning-based methods. General information of these tools, including their underlying approaches and training datasets, is summarized in **Table** 3.

**Table 3.** General information of protein stability prediction models used for benchmarking.

| Tool | Method | Training Set |
| --- | --- | --- |
| ThermoMPNN (10) | Graph neural network | Megascale + FireProt |
| ThermoMPNN-D (11) | Graph neural network | Megascale |
| SPURS (18) | Protein language model and graph neural network | Megascale |
| PROSTATA (39) | Transformer model | S2648 + Q3421 + VariBench |
| DDGEmb (30) | Transformer model | S2450 |
| DDMut (44) | Graph neural network | S9028 |
| DDGun-3D (23) | Structure-based | – |
| DDGun-seq (23) | Sequence-based | – |
| Schrödinger Maestro (31) | Force field-based scoring method | – |
| MOE (14) | Force field-based scoring method | – |
| Rosetta (17, 34) | Empirical/statistical potential method | – |
| FoldX (9) | Empirical/statistical potential method | – |

When evaluated on the full curated enzyme mutation dataset **Table** 4, ML-based models consistently outperformed non-ML approaches across correlation and error metrics. Among all models, PROSTATA achieved the best overall performance, exhibiting the highest PCC (0.792) and SCC (0.866), together with the lowest RMSE (1.194). ThermoFusion also demonstrated strong predictive ability (PCC = 0.668, SCC = 0.668, RMSE = 1.522), outperforming several existing ML models. ThermoMPNN, ThermoMPNN-D, Spurs, and DDGEmb showed moderate but consistent performance. In comparison, non-ML tools generally exhibited weaker linear correlations and higher prediction errors. For example, Maestro showed almost no linear correlation (PCC = 0.080) despite a moderate rank correlation (SCC = 0.554), indicating limited predictive reliability. MOE and DDGun_Seq displayed modest correlations across both metrics, suggesting only moderate predictive performance. While Rosetta achieved relatively strong correlations (PCC = 0.557, SCC = 0.630), it was associated with the highest prediction error (RMSE = 6.176) among the tested tools, indicating extreme outliers in the predictions. Similarly, FoldX and DDGun_3D reached moderate correlation levels but still showed comparatively high RMSE values, exhibiting a trade-off between ranking ability and absolute prediction accuracy. It is important to note that for force-field-based methods, such as Schrödinger Maestro and MOE, performance is highly sensitive to the choice of sampling parameters, making direct comparison with ML-based approaches inherently challenging. Additionally, although Rosetta demonstrates comparatively better correlation than other non-ML methods, it is substantially more computationally intensive and time-consuming, which may limit its practical applicability.

**Table 4.** Benchmarking Protein Mutation Thermostability Prediction Tools on the Curated Enzyme Dataset (All Mutations)

| ML Models |  |  |  |
| --- | --- | --- | --- |
| Method | PCC | SCC | RMSE |
| ThermoMPNN | 0.601 | 0.642 | 1.663 |
| ThermoMPNN-D | 0.588 | 0.634 | 1.706 |
| SPURS | 0.604 | 0.634 | 1.717 |
| PROSTATA | **0.792** | **0.866** | **1.194** |
| DDMut | 0.656 | 0.724 | 1.570 |
| DDGEmb | 0.617 | 0.745 | 1.545 |
| ThermoFusion | 0.668 | 0.668 | 1.522 |
| Non-ML Models |  |  |  |
| Method | PCC | SCC | RMSE |
| MAESTRO | 0.080 | 0.554 | 2.010 |
| MOE | 0.431 | 0.466 | 1.925 |
| Rosetta | 0.557 | 0.630 | 6.176 |
| FoldX | 0.467 | 0.588 | 2.559 |
| DDGun-3D | 0.515 | 0.474 | 1.842 |
| DDGun-Seq | 0.454 | 0.454 | 1.911 |

To further dissect model behavior, we evaluated performance across three mutation regimens: destabilizing mutations (ΔΔ*G <* − 1.0 kcal/mol), neutral mutations ( − 1.0 kcal/mol ≤ ΔΔ*G* ≤ 1.0 kcal/mol), stabilizing mutations (ΔΔ*G >* 1.0 kcal/mol). The distribution of mutation ΔΔ*G* values of the curated enzyme dataset is shown in **Figure** 2 (left), and the prediction analysis results are shown in **Table** 5. A consistent trend exhibited across all methods: both ML and non-ML tools performed substantially better in predicting destabilizing mutations compared to neutral and stabilizing ones. In this regime, most models achieved their highest correlation coefficients, with several ML-based approaches (e.g., PROSTATA, DDMut, and ThermoFusion) showing moderate to strong agreement with experimental values. Among them, PROSTATA reached the best overall performance (PCC = 0.792, SCC = 0.866), while even non-ML methods such as Rosetta and FoldX achieved their strongest correlations in this subset. This suggests that large destabilizing effects are more readily captured by current predictive frameworks. In contrast, prediction performance decreased noticeably for neutral mutations. Across both ML and non-ML methods, correlation coefficients were consistently lower, reflecting the difficulty of resolving subtle stability changes within this ΔΔ*G* region. Although some models (e.g., PROSTATA and DDGemb) retained moderate rank correlations, overall predictive accuracy remained limited, with reduced sensitivity to the neutral mutation effects. The most pronounced performance degradation was observed for stabilizing mutations. In this regime, most models exhibited weak or even negative PCC results, indicating a systematic inability to correctly capture stabilizing effects. This trend was consistent across both ML-based and physics-based approaches, suggesting that stabilizing mutations remain particularly challenging to model. Even when rank correlations were marginally positive in some cases, the overall predictive reliability was poor, accompanied by high RMSE values. Overall, these results reveal a fundamental asymmetry in current stability prediction methods: destabilizing mutations are captured more reliably, whereas neutral and especially stabilizing mutations remain challenging to predict. This is particularly relevant for enzyme engineering, where identifying stabilizing variants is often a primary objective.

**Fig. 2.**
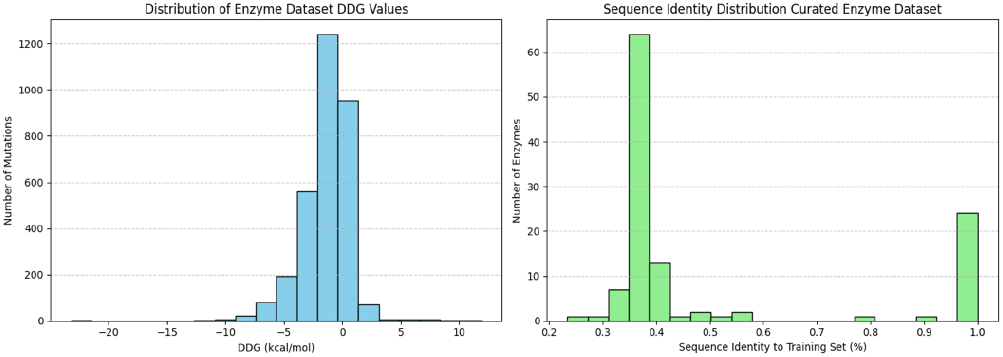
ΔΔ*G* (left) and Sequence Identity (right) Distributions of the Curated Enzyme Dataset.

**Table 5.**
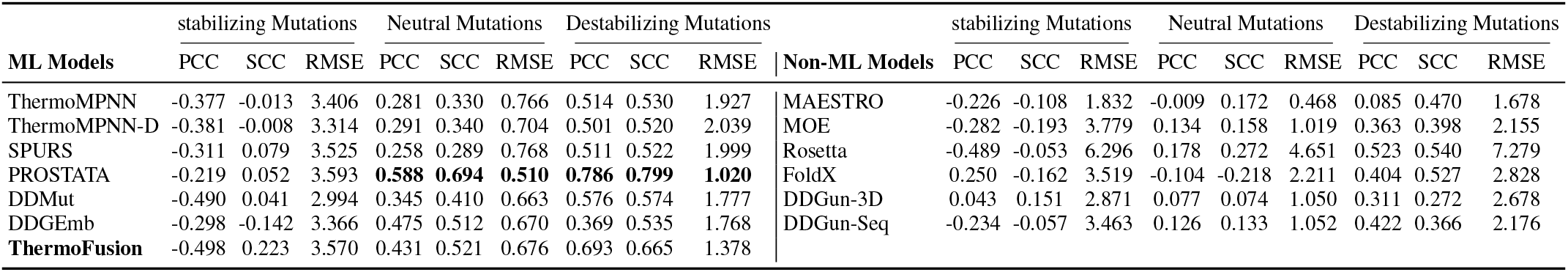
Benchmarking Protein Mutation Thermostability Prediction Tools on the Curated Enzyme Dataset Based on Mutation Types.

| ML Models | stabilizing Mutations |  |  | Neutral Mutations |  |  | Destabilizing Mutations |  |  | Non-ML Models | stabilizing Mutations |  |  | Neutral Mutations |  |  | Destabilizing Mutations |  |  |
| --- | --- | --- | --- | --- | --- | --- | --- | --- | --- | --- | --- | --- | --- | --- | --- | --- | --- | --- | --- |
|  | PCC | SCC | RMSE | PCC | SCC | RMSE | PCC | SCC | RMSE |  | PCC | SCC | RMSE | PCC | SCC | RMSE | PCC | SCC | RMSE |
| ThermoMPNN | -0.377 | -0.013 | 3.406 | 0.281 | 0.330 | 0.766 | 0.514 | 0.530 | 1.927 | MAESTRO | -0.226 | -0.108 | 1.832 | -0.009 | 0.172 | 0.468 | 0.085 | 0.470 | 1.678 |
| ThermoMPNN-D | -0.381 | -0.008 | 3.314 | 0.291 | 0.340 | 0.704 | 0.501 | 0.520 | 2.039 | MOE | -0.282 | -0.193 | 3.779 | 0.134 | 0.158 | 1.019 | 0.363 | 0.398 | 2.155 |
| SPURS | -0.311 | 0.079 | 3.525 | 0.258 | 0.289 | 0.768 | 0.511 | 0.522 | 1.999 | Rosetta | -0.489 | -0.053 | 6.296 | 0.178 | 0.272 | 4.651 | 0.523 | 0.540 | 7.279 |
| PROSTATA | -0.219 | 0.052 | 3.593 | **0.588** | **0.694** | **0.510** | **0.786** | **0.799** | **1.020** | FoldX | 0.250 | -0.162 | 3.519 | -0.104 | -0.218 | 2.211 | 0.404 | 0.527 | 2.828 |
| DDMut | -0.490 | 0.041 | 2.994 | 0.345 | 0.410 | 0.663 | 0.576 | 0.574 | 1.777 | DDGun-3D | 0.043 | 0.151 | 2.871 | 0.077 | 0.074 | 1.050 | 0.311 | 0.272 | 2.678 |
| DDGEmb | -0.298 | -0.142 | 3.366 | 0.475 | 0.512 | 0.670 | 0.369 | 0.535 | 1.768 | DDGun-Seq | -0.234 | -0.057 | 3.463 | 0.126 | 0.133 | 1.052 | 0.422 | 0.366 | 2.176 |
| **ThermoFusion** | -0.498 | 0.223 | 3.570 | 0.431 | 0.521 | 0.676 | 0.693 | 0.665 | 1.378 |  |  |  |  |  |  |  |  |  |  |

The observed bias toward detecting destabilization may therefore restrict the discovery of beneficial variants and lead to underestimation of stabilizing effects. These prediction performances may partially reflect biases in the training data, such as an imbalance in the representations of mutation types. However, it is more likely rooted in the methodological limitations common to both ML and physics-based approaches, particularly those relying on static structural representations with limited treatment of solvation and conformational dynamics. In this context, approaches that explicitly account for protein flexibility and solvent effects – such as molecular dynamics-based methods and alchemical free energy calculations (e.g., FEP+) – may provide improved accuracy, especially for neutral and stabilizing mutations where subtle energetic contributions play a critical role.

### D. Evaluation of Model Generalization Beyond Training Distribution

To assess model generalization, curated enzymes were clustered by sequence identity relative to each model’s training set ( ≤ 40% or *>* 40%) using CD-HIT. This analysis was limited to ML-based models, since non-ML tools do not disclose their training data. Most models showed decreased predictive performance for enzymes with ≤ 40% sequence identity (**Table** 6), including pronounced declines in correlation for PROSTATA, DDMut, DDGemb, and moderate PCC values (0.573 and 0.498) for ThermoMPNN and ThermoMPNN-D. ThermoFusion, however, maintained strong performance (PCC = 0.653 and SCC = 0.653), suggesting improved robustness to evolutionary distance. This observation consistently shows that ESM2 embeddings provide evolutionary and functional context. By integrating both modalities, ThermoFusion generalizes effectively to evolutionarily remote enzymes and captures stability-determining factors that go beyond simple memorization. These findings align with recent advances in protein-ligand binding affinity predictions where models combining structural and contextual embeddings outperform only structure- or sequence-based methods. Overall, this multi-modal architecture enables ThermoFusion to learn causal determinants of stability rather than relying on sequence and structure memorization.

**Table 6.** Prediction performance of ML-based models on enzymes sharing *≤*40% sequence identity with their corresponding training sets.

| Model | PCC | SCC | RMSE |
| --- | --- | --- | --- |
| ThermoMPNN | 0.573 | 0.576 | 1.733 |
| ThermoMPNN-D | 0.498 | 0.555 | 1.933 |
| SPURS | 0.435 | 0.551 | 1.821 |
| PROSTATA | 0.373 | 0.488 | 2.036 |
| DDMut | 0.363 | 0.463 | 2.093 |
| DDGEmb | 0.315 | 0.504 | 1.954 |
| **ThermoFusion** | **0.653** | **0.653** | **1.551** |

When sequence identity exceeded 40% (see Supplementary Table S2), predictive correlations increased and RMSE decreased significantly across all machine learning models. The sequence identity distribution of the curated enzyme dataset is shown in **Figure** 2 (right).

## Conclusions

In this study, we developed ThermoFusion, a hybrid deep learning framework that integrates both sequence- and structure-based information for predicting the effects of single-point mutations on protein stability (ΔΔ*G*). By combining structural embeddings from ThermoMPNN with sequence embeddings from the pretrained protein language model ESM2, ThermoFusion effectively models the mutations that influence protein thermostability.

The comparison with the baseline model, ThermoMPNN, indicated that correctly incorporating ESM2 sequence embeddings provided complementary evolutionary context. The ThermoFusion model, which integrates both sequence and structural representations, consistently outperformed the baseline method across five benchmark datasets. These results highlight the synergistic effect of integrating sequence and structural information, particularly for proteins with diverse sequences and structures.

When applied to a curated enzyme mutation dataset comprising 105 enzymes and 3144 mutation sites, ThermoFusion showed strong predictive performance, outperforming several existing ML and non-ML methods in correlation and error metrics. Notably, ThermoFusion maintained robust performance on out-of-distribution enzymes with low sequence identity ( ≤ 40%) to the training set, indicating improved generalization to evolutionary distant proteins. By contrast, several other ML-based prediction models exhibited substantial performance degradation in this low-identity regime, suggesting a stronger dependence on similarity to sequences seen during training. Such dependence on sequence similarity can lead to data leakage effects, where training and validation datasets contain closely related proteins or mutations. In these scenarios, models may achieve artificially high benchmark performance by memorizing patterns associated with similar sequence rather than learning the underlying factors of protein stability.

Ensuring strong generalization is particularly important in enzyme engineering, where prediction models are often applied to proteins that differ substantially from those used during model training. In real-world applications, researchers frequently seek to engineer enzymes from new organisms, alter substrate specificity, or improve stability under industrial conditions such as elevated temperatures, extreme pH, or organic solvents. These tasks commonly involve exploring sequence space that lies far from existing experimental dataset.

Another important observation from our analysis is the systematic variation in predictive performance across mutation regimes. Most evaluated models performed better on strongly destabilizing mutations, while prediction performance decreased significantly for neutral and stabilizing mutations. This suggests that current methods are more sensitive to large energetic shifts than to stabilizing changes. This may be partly influenced by training dataset composition, including imbalances across mutation types and relative fewer well-characterized stabilizing variants in the training data. However, it could more fundamentally reflect the challenges associated with modeling small free-energy differences using methods that rely primarily on static structural representations and limited treatment of conformational flexibility and solvation effects. Improving performance across all mutation regimes will likely require both more balanced and diverse training datasets and methodological advances that better capture protein dynamics.

Overall, the practical implications of ThermoFusion are significant for rational enzyme engineering. Accurate prediction of mutation-induced stability changes can significantly reduce the experimental burden associated with large-scale mutant library screening, guide the design of thermostable enzymes for industrial and biomedical applications, and accelerate the development of proteins with desired functional properties. Furthermore, the multi-modal design of ThermoFusion provides a flexible framework that can be extended to broader protein engineering tasks, including multi-point mutation optimization, functional variant prioritization, and structure-guided protein design. Future work will focus on improving prediction accuracy for neutral and stabilizing mutations and further enhancing the model’s generalization capability across diverse protein families and engineering studies.

## Methods

### E. Structural Embedding via ThermoMPNN

Protein structure information is encoded using a pretrained Protein-MPNN encoder integrated within the ThermoMPNN framework. Given a protein backbone comprising *L* residues, the structure is represented by backbone atomic coordinates 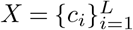, where 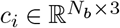 denotes the coordinates of the backbone atoms for residue *i*, together with the residue sequence *S*.

ProteinMPNN represents the protein as a graph in which residues interact through spatially neighboring connections derived from the 3D backbone structure. Message passing operations are performed across the graph to iteratively update residue representations based on local geometric context and neighboring residues.

Let the hidden representations from the final *K* encoder layers be denoted as 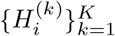, where *K* corresponds to the number of retained final layers. These hidden representations are concatenated with the ProteinMPNN residue embedding 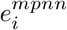 to form the final structural feature vector:

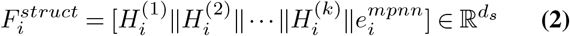

where ‖ denotes vector concatenation and *d*_*s*_ is the dimensionality of the structural representation. These embeddings encode residue environment, backbone geometry, chain context, and long-range structural dependencies learned through graph message passing.

### F. Sequence Embedding via ESM2

To capture sequence-derived functional and evolutionary information, the amino acid sequence *A* = (*a*_1_, …, *a*_*L*_) is encoded using the pretrained ESM2 (esm2_t33_650M_UR500) model. ESM2 generates contextualized residue-level embedding by modeling long-range dependencies and evolutionary constraints directly from large-scale sequence data. The resulting sequence representation for residue *i* is defined as:

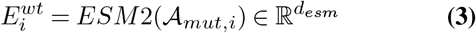

where *d*_*esm*_ denotes the dimensionality of the ESM2 model. These embeddings capture sequence-derived features, mutational sensitivity, and global context relevant to protein stability.

### G. Structure-Preserved Conditional Fusion

To integrate structural information into the ESM2 embedding space, ProteinMPNN-derived structural features are projected through a nonlinear modulation network:

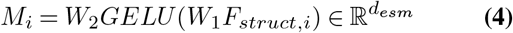

where *W*_1_ and *W*_2_ are learnable linear projections. Layer normalization and dropout regularization are applied within the modulation network.

A learnable gating mechanism controls the influence of structural conditioning:

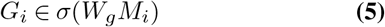

where, 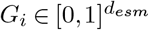 . The fused residue representations are computed as:

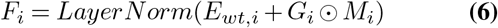

where, ⊙ denotes element-wise multiplication. This gated fusion mechanism enables adaptive integration of structural conditioning with sequence-derived semantic information.

### H. Mutation Effect Encoding

To explicitly model mutation-induced sequence perturbation, the embedding difference between mutation and wild-type residues is computed:

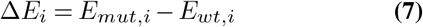

where, *E*_*mut,i*_ = *ESM* 2( *A*_*mut,i*_). This perturbation vector is passed through a multilayer reduction network consisting of stacked linear layers, ReLU activations, and dropout regulation:

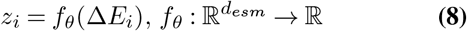

This produces a scalar mutation effect score: *z*_*i*_ ∈ ℝ.

## I. Final ΔΔ*G* Prediction

For each mutated residue, the fused structure-conditioned representation *F*_*i*_ is concatenated with the scalar mutation effect score *z*_*i*_:

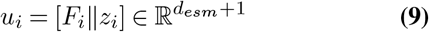

The resulting feature vector is passed through a MLP regression:

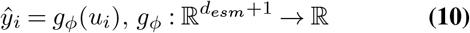

The predicted values: 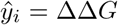 corresponds to the estimated thermostability change for the mutation.

### J. Model Training

The ThermoFusion model was first pre-trained on the Megascale dataset (38), which contains 272,712 ΔΔ*G* measurements spanning 331 natural and 148 *de novo* designed protein domains, each 40 - 72 residues in length. The model was subsequently fine-tuned on the FireProt dataset (36), comprising of 3,438 mutations across 100 unique proteins. Both datasets, curated by Dieckhaus et al. (10), were split into training, validation, and test sets using an 80/10/10 ratio. For each mutation, the model generated a predicted ΔΔ*G* value. Predictions were compared with experimentally measured ΔΔ*G* values using the Huber loss:

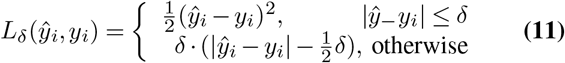

where 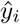 is the predicted stability change and *y*_*i*_ is the experimental value. A Huber loss with *δ* = 1.0 was used to reduce sensitivity to extreme outliers. Predicted values were clipped to the range [ −15, 15] before loss calculation and metric evaluation.

Models were pre-trained on Megascale dataset for 20 epochs and subsequently fine-tuned on the FireProt dataset for 50 epochs. Further increases in model training duration did not improve prediction performance. During training model performance was monitored using Huber loss together with *R*^2^, mean square error (MSE), root mean squared error (RMSE), and Spearman correlation coefficient (SCC) calculated against experimental ΔΔ*G* values.

### K. Evaluation Metrics

To evaluate model performance, we employed three widely used metrics to assess the prediction results of different methods: root mean square error (RMSE), Spearman’s correlation coefficient (SCC), and Pearson’s correlation coefficient (PCC) (10, 30).

RMSE is a commonly used metric that measures the difference between predicted and actual values. Its core idea is to quantify the magnitude of prediction errors and provide a single numerical value that reflects the model’s overall accuracy. It is defined as:

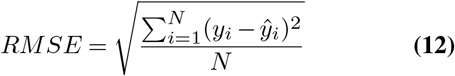

where *N* is the total number of observations; *y*_*i*_ denotes the experimental mutation ΔΔ*G* value; 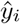 denotes the predicted ΔΔ*G* value.

PCC measures the linear correlation between two set of variables. Its value ranges from -1 to 1, where values closer to -1 to 1 indicate stronger negative or positive linear correlations, respectively. PCC is calculated as:

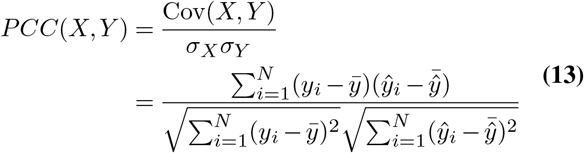

where *COV* (*X, Y* ) is the covariance between the two variables, and *σ*_*X*_ and *σ*_*Y*_ are their respective standard deviations. SCC evaluates the strength of a monotonic relationship between two variables, regardless of whether the relationship is linear or non-linear. The calculation of SCC is based on the ranks of the variables rather than their raw values, and is given by:

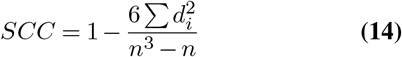

where *d*_*i*_ is the difference between the ranks of the two variables for the *i*-th observation, and *n* is the total number of observations.

### L. Enzyme Dataset Curation

To evaluate ThermoFusion’s performance on enzymes, we curated a dataset of single-point enzyme mutations from the five datasets used in the baseline comparison study. The enzyme data were collected from the FireProt, S669, Q3421, and ThermoMutDB datasets. During data curation, each protein PDB ID in these datasets was manually verified in the Protein Data Bank to ensure that only enzymes were retained. We then removed mutations that were not single-point mutation, deleted duplicate enzymes and mutation entries, and excluded mutations lacking recording ΔΔ*G* values. Due to missing residues in crystallographic PDB structures, mutation positions were aligned to the corresponding structures using a Python script and subsequently verified manually. For ΔΔ*G* values, ΔΔ*G <* 0.0 kcal/mol indicates mutations with destabilizing effects on protein folding, whereas ΔΔ*G >* 0.0 kcal/mol indicates mutations with stabilizing effects on protein folding.

### M. Model Implementation

The machine learning models were developed in Python (40). Deep learning methods were implemented using PyTorch (28). ThermoMPNN protein structure embeddings were extracted using the ProteinMPNN v_48_020 model weights (1). Protein sequence embeddings were obtained via ESM2 (esm2_t33_650M_UR500) (22) with 1280-dimensional token representations from the last layer.

## Data Availability

The ThermoFusion model code is available on GitHub: https://github.com/yaow1004/ThermoFusion. The trained model weights, datasets, and benchmarking results are available on zenodo: https://zenodo.org/records/20683630. The materials are additionally archived in the University of Milan Dataverse (institutional FAIR repository), where applicable, at the following link: https://doi.org/10.13130/RD_UNIMI/TJDYNW.

## Supporting Information

**Table S1. Curated Enzyme Dataset information.**

**Table S2. Prediction Performance of ML-Based Models on Enzyme Sharing *>* 40% Sequence Identity with Their Corresponding Training Sets.**

**Text 1. Schrödinger Maestro and MOE Residue Scanning Methods**.

## Acknowledgments

This project has received funding from the European Union’s Horizon Europe research and innovation programme under the Marie Skłodowska-Curie grant agreement No. 101073546 (MSCA Doctoral Network Metal-containing Radical Enzymes - MetRaZymes), and grants from MUR - “Progetto Eccellenza 2023 - 2027”.

Computational resources provided by INDACO Core facility, which is a project of High-Performance Computing at the University of Milan https://www.unimi.it.

Grateful thanks to Dr. Florian Kleinbeck, Dr. Frederic Stanger and the Development Chemical and Analytical Department of Novartis Pharma AG environment for hosting and organizing the PhD industrial secondment. We also thank Dr. Maria Laura De Sciscio, for discussing and providing suggestions for the manuscript.

## Author Contributions

**Conceptualization:** Fabian Meyer

**Data curation:** Yao Wei

**Formal analysis:** Yao Wei

**Funding acquisition:** Ivano Eberini

**Investigation:** Yao Wei, Fabian Meyer

**Methodology:** Yao Wei, Fabian Meyer

**Project administration:** Ivano Eberini

**Resources:** Fabian Meyer, Ivano Eberini

**Software:** Fabian Meyer, Ivano Eberini

**Supervision:** Fabian Meyer

**Validation:** Yao Wei, Fabian Meyer

**Visualization:** Yao Wei

**Writing - original draft:** Yao Wei

**Writing - review & editing:** Yao Wei, Fabian Meyer, Ivano Eberini

