## Supplementary Material for "ThermoFusion: A Multimodal Deep Learning Framework for Generalizable Prediction of Enzyme Thermostability"

### Table S1. Curated Enzyme Dataset Information

| PDB | Enzyme name | Enzyme class |
|---|---|---|
| 1AMQ | Pyridoxamine 5'-phosphate-type aspartate aminotransferases | Transferase |
| 1B26 | Glutamate dehydrogenase | Oxidoreductase |
| 1BLC | Inhibition of beta-lactamase by Clavulanate | Hydrolase |
| 1DIL | Sialidase from salmonella typhimurium complexed with apana and epana inhibitors | Hydrolase |
| 1DPM | Zinc-containing phosphotriesterase with bound substrate analog diethyl 4-methylbenzylphosphonate | Hydrolase |
| 1FKJ | FKBP12-FK506, an immunophilin immunosuppressant complex | Isomerase |
| 1HTI | Recombinant human triosephosphate isomerase | Isomerase |
| 1IDS | Iron-dependent superoxide dismutase from mycobacterium tuberculosis | Oxidoreductase |
| 1LUC | Bacterial luciferase | Oxidoreductase |
| 1N0J | Human Mitochondrial MN3+ Superoxide Dismutase | Oxidoreductase |
| 1OH0 | Ketosteroid isomerase complexed with equilenin | Isomerase |
| 1P2P | Porcine pancreatic phospholipase | Hydrolase |
| 1PIN | PIN1 peptidyl-prolyl cis-trans isomerase from homo sapiens | Isomerase |
| 1QM4 | Methionine Adenosyltransferase Complexed with a L-Methionine Analogue | Transferase |
| 1RGG | Hydrolase, Guanyloribonuclease | Hydrolase |
| 1SHF | Comparison of 3D structures of sh3 domains in tyrosine kinases and spectrin | Transferase |
| 2CRK | Muscle creatine kinase | Transferase |
| 2TS1 | Interaction of tyrosyl-t/rna synthetase with the tyrosyl adenylate intermediate | Ligase |
| 2WSY | Wild-type tryptophan synthase | Lyase |
| 1BNI | Barnase wildtype structure at pH 6.0 | Hydrolase |
| 1EY0 | Wild-type S.nuclease | Hydrolase |
| 1G4I | Bovine pancreatic phospholipase | Hydrolase |
| 1HMK | Recombinant goat alpha-lactalbumin | Transferase |
| 1LNI | A ribonuclease from streptomyces aureofaciens | Hydrolase |
| 1LZ1 | Human lysozyme | Hydrolase |
| 1MGR | RNase Sa3, cytotoxic microbial ribonuclease | Hydrolase |
| 1ONC | P-30, an amphibian ribonuclease with anti-tumor activity | Hydrolase |
| 1PDO | Phosphoenolpyruvate-dependent phosphotransferase system | Transferase |
| 1SUP | Subtilisin bpn' | Hydrolase |
| 1TTQ | Tryptophan synthase in the presence of potassium | Lyase |
| 1WQ5 | Tryptophan synthase alpha-subunit from E. coli | Lyase |
| 1ZG4 | TEM1 beta lactamase | Lyase |
| 2LZM | Bacteriophage t4 lysozyme | Hydrolase |
| 2RN2 | Ribonuclease h from E. coli | Hydrolase |
| 3ECA | E. coli L-asparaginase, an enzyme used in cancer therapy (elspar) | Hydrolase |
| 3GLY | Glucoamylase from aspergillus awamori var. x100 | Hydrolase |
| 3PGK | Yeast phosphoglycerate kinase | Transferase |
| 4LYZ | Hen egg-white lysozyme | Hydrolase |
| 5DFG | C-As lyase with mutations Y100H and V102F | Lyase |
| 1ANK | E. coli adenylate kinase with bound AMP and AMPPNP | Transferase |
| 1CHK | Streptomyces n174 chitosanase ph5.5 298k | Hydrolase |
| 1IO2 | Type 2 ribonuclease h from hyperthermophilic archaeon, thermococcus kodakaraensis kod1 | Hydrolase |
| 1KE4 | AmpC beta-lactamase from E. coli | Hydrolase |
| 1KFW | Catalytic domain of psychrophilic chitinase B from Arthrobacter TAD20 | Hydrolase |
| 1L63 | Analysis of the interaction between charged side chains and the alpha-helix dipole using designed thermostable mutants of phage t4 lysozyme | Hydrolase |
| 1RN1 | GLN 25-ribonuclease t1 at 1.84 angstroms resolution | Hydrolase |
| 3SIL | Sialidase from salmonella typhimurium | Hydrolase |
| 4BLM | Beta-lactamase of bacillus licheniformis 749(slash)c | Hydrolase |
| 1A23 | Reduced dsba from e. coli | Oxidoreductase |
| 1APS | Cylphosphatase | Hydrolase |

Continued on next page

**Table S1. Curated Enzyme Dataset Information continued**

| PDB | Enzyme name | Enzyme class |
|---|---|---|
| 1BA3 | Firefly luciferase in complex with bromoform | Oxidoreductase |
| 1BKS | Tryptophan synthase from salmonella typhimurium | Lyase |
| 1BP2 | Bovine pancreatic phospholipase a2 | Hydrolase |
| 1BU4 | Ribonuclease 1 complex with 2'-GMP | Hydrolase |
| 1CAH | Cobalt carbonic anhydrase complexed with bicarbonate | Lyase |
| 1CM2 | HIS15ASP HPR after hydrolysis of ringed species | Transferase |
| 1AM7 | Lysozyme from bacteriophage lambda | Hydrolase |
| 1D5G | PDZ2 domain from human phosphatase HPTP1E complexed with a peptide | Hydrolase |
| 1DFO | E. coli serine hydroxymethyltransferase in complex with glycine and 5-formyl tetrahydrofolate | Transferase |
| 1EL1 | Canine milk lysozyme (holo-type) | Hydrolase |
| 1EW6 | Dehaloperoxidase from amphitrite ornata indicate common ancestry with globins | Oxidoreductase |
| 1GLM | Glucoamylase from aspergillus awamori var. x100 | Hydrolase |
| 1JLV | Anopheles dirus species B glutathione S-transferases 1-3 | Transferase |
| 1N18 | Thermostable mutant of Human Superoxide Dismutase, C6A, C111S | Oxidoreductase |
| 1NEY | Triosephosphate Isomerase in Complex with DHAP | Isomerase |
| 1NMV | human Pin1 | Isomerase |
| 1O6X | Activation domain of human procarboxypeptidase A2 | Hydrolase |
| 1OSI | 3-Isopropylmalate dehydrogenase | Oxidoreductase |
| 1QGD | Transketolase from E. coli | Transferase |
| 1RIL | Ribonuclease H from thermus thermophilus HB8 | Hydrolase |
| 1RTB | Thymidylic acid tetramer in complex with ribonuclease A | Hydrolase |
| 1RX4 | Dihydrofolate reductase complexed with 5,10-dideazatetrahydrofolate and 2'-monophosphoadenosine 5'-diphosphoribose | Oxidoreductase |
| 1SPD | Amyotrophic lateral sclerosis and structural defects in Cu, Zn superoxide dismutase | Oxidoreductase |
| 1STN | Staphylococcal nuclease | Hydrolase |
| 1THQ | Outer Membrane Enzyme PagP | Oxidoreductase |
| 1W4H | Peripheral-subunit from mesophilic, thermophilic and hyperthermophilic bacteria fold by ultrafast, apparently two-state transitions | Transferase |
| 1WPQ | Ternary Complex of Glycerol 3-phosphate Dehydrogenase 1 with NAD and dihydroxyactone | Oxidoreductase |
| 1XWS | Human PIM1 kinase domain | Transferase |
| 1XXN | A mesophilic xylanase A from Bacillus subtilis 1A1 | Hydrolase |
| 1YPI | Yeast triosephosphate isomerase | Isomerase |
| 1Z1I | native SARS CLpro | Hydrolase |
| 2ADA | Adenosine deaminase complexed with a transition-state analog | Hydrolase |
| 2AKY | Adenylate kinase from yeast ligated with inhibitor AP5a, showing the pathway of phosphoryl transfer | Transferase |
| 2BJD | Sulfolobus Solfataricus Acylphosphatase. Triclinic space group | Hydrolase |
| 2EWN | E. coli Biotin Repressor with co-repressor analog | Ligase |
| 2JIE | Beta-glucosidase B from bacillus polymyxa complexed with 2-F-glucose | Hydrolase |
| 2JYS | Simian Foamy Virus (mac) protease | Hydrolase |
| 2K7J | Human Acylphosphatase (AcPh) surface charge-optimized | Hydrolase |
| 2VY0 | Endo-beta-1,3-glucanase from Pyrococcus furiosus | Hydrolase |
| 2WZB | Human phosphoglycerate kinase in complex with ADP, 3PG and magnesium trifluoride | Transferase |
| 3BDC | Staphylococcal nuclease variant Delta+PHS at cryogenic temperature | Hydrolase |
| 3BLS | AMPC beta-lactamase from E. coli | Hydrolase |
| 3D2A | A thermostable mutant of Bacillus subtilis lipase | Hydrolase |
| 3ECU | Human apo Cu,Zn Superoxide Dismutase (SOD1) | Oxidoreductase |
| 3GDL | Orotidine 5'-monophosphate decarboxylase from Saccharomyces cerevisiae complexed with 6-azauridine 5'-monophosphate | Lyase |
| 4Q0M | Pyrococcus furiosus L-asparaginase | Hydrolase |
| 5CG0 | Spodoptera frugiperda Beta-glycosidase | Hydrolase |

Continued on next page

#### Table S1. Curated Enzyme Dataset Information continued

| PDB | Enzyme name | Enzyme class |
|---|---|---|
| 5NP8 | PGK1 in complex with CRT0063465 (3-[2-(4-bromophenyl)-5,7-dimethyl-pyrazolo[1,5-a]pyrimidin-6-yl]propanoic acid) | Transferase |
| 5TR5 | Serine 65 phosphorylated UBL domain from parkin | Ligase |
| 1F8I | Isocitrate lyase:nitropropionate:glyoxylate complex from mycobacterium tuberculosis | Lyase |
| 1IR3 | Phosphorylated insulin receptor tyrosine kinase in complex with peptide substrate and ATP analog | Transferase |
| 2ARF | Wilson ATPase N-domain in the presence of ATP | Hydrolase |
| 2N7Z | RIP2 CARD | Transferase |
| 3BCI | Staphylococcus aureus DsbA | Oxidoreductase |
| 1AQH | Alpha-amylase from alteromonas haloplanctis | Hydrolase |
| 1BCX | Bacillus circulans xylanase | Hydrolase |

**Table S2. Prediction Performance of ML-Based Models on Enzymes Sharing $>$40% Sequence Identity with Their Corresponding Training Sets**

| ML Model | PCC | SCC | RMSE |
|---|---|---|---|
| ThermoMPNN | 0.625 | 0.696 | 1.600 |
| ThermoMPNN-D | 0.683 | 0.707 | 1.465 |
| SPURS | 0.714 | 0.695 | 1.635 |
| PROSTATA | 0.842 | 0.902 | 1.032 |
| DDMut | 0.737 | 0.778 | 1.423 |
| DDGEmb | 0.652 | 0.775 | 1.496 |
| **ThermoFusion** | **0.808** | **0.876** | **1.172** |

#### 1. Schrödinger Maestro and MOE Residue Scanning Methods

Before performing residue scanning, all enzymes underwent protein structure preparation using the corresponding software. Residue scanning calculations were then conducted with two computational programs to predict the thermostability changes in the curated enzyme dataset. BioLuminate in the Schrödinger Maestro Suite (2025-1) employed side-chain prediction with backbone minimization and a cut-off of 0.00 Å to assess the relative structural stability of the mutants. For residue scanning in Molecular Operating Environment (MOE, 2024.06), the Amber:EHT force field with R-Field solvation was applied. The LowMode ensemble method was selected with a maximum of 50 iterations, a 0.25 Å RMSD limit, and up to 25 conformations.

All benchmarking work on the licensed prediction tools, including Schrödinger Maestro, MOE, Rosetta, and FoldX, was conducted at the University of Milan.
